# First Indian report on Genome-wide Comparison of Multidrug-Resistant *Escherichia coli* from Blood Stream Infections

**DOI:** 10.1101/705905

**Authors:** Naveen Kumar Devanga Ragupathi, Balaji Veeraraghavan, Dhiviya Prabaa Muthuirulandi Sethuvel, Shalini Anandan, Karthick Vasudevan, Ayyan Raj Neeravi, Jones Lionel Kumar Daniel, Sowmya Sathyendra, Ramya Iyadurai, Ankur Mutreja

## Abstract

**Background:** Multidrug-resistant (MDR) *E. coli* with extended-spectrum β-lactamases (ESBLs) is becoming endemic in health care settings around the world. Baseline data on virulence and AMR of specific lineages of *E. coli* circulating in developing countries like India is currently lacking.

**Methods:** Whole-genome sequencing was performed for 60 MDR *E. coli* isolates. Genome-wide analysis was performed at single nucleotide polymorphism (SNP) level resolution to identify the relation between the isolates in context of time, virulence and AMR determinants possessed.

**Results:** Genome comparison revealed the presence of ST-131 global MDR and ST410 as emerging-MDR clades of *E. coli*. AMR gene profile for cephalosporin and carbapenem resistance differed between the clades. Genotypes *bla*_CTX-M-15_ and *bla*_NDM-5_ were common among cephalosporinases and carbapenemases, respectively. For aminoglycoside resistance, *rmtB* was positive for 31.7% of the isolates, of which 30% were co-harboring carbapenemases. Further, the FimH types and virulence gene profile positively correlated with the SNP based phylogeny, which also revealed the evolution of MDR clones among the study population with temporal accumulation of SNPs. The predominant clone was ST167 (*bla*_NDM_ lineage) followed by ST405 (global clone ST131 equivalent) and ST410 (fast spreading high risk clone).

**Conclusions:** This is the first report on the whole genome analysis of MDR *E. coli* lineages circulating in India. Data from this study will provide public health agencies a baseline portfolio of AMR and virulence in pathogenic *E. coli* in the region.

## Introduction

*Escherichia coli* is the leading cause of bloodstream infections (BSIs) caused by Gram-negative bacteria [1] and other common infections including urinary tract infections (UTIs). As an important commensal component of the biosphere, *E. coli* is colonizes the lower gut of animals and humans and gets released in the environment as a norm allowing widespread dissemination.

Virulence of *E. coli* is driven by multiple factors that involves adhesins, toxins, siderophores, lipopolysaccharide (LPS), capsule, and invasins [2]. It was reported that a large proportion of MDR *E. coli* carried by people are acquired via food, especially from farm animals [3]. Although most of the MDR *E. coli* are reported to be community acquired rather than healthcare [4,5], recently MDR *E. coli*, which produce extended-spectrum β-lactamases (ESBL) have been endemic in health care settings.

Among MDR *E. coli*, AMR mediated by ESBL is mainly due to the *bla*CTX-M family, particularly *bla*CTX-M-15 and 14, compared to the less frequent observation of *bla*SHV and *bla*OXA families [6–8]. Carbapenem resistance in *E. coli* was mostly reported to be mediated by *bla*OXA-48 [9], *bla*NDM and *bla*VIM [10]. Also, AMR among *E. coli* is increasingly reported for fluoroquinolones and third- and fourth-generation cephalosporins. Sequence type 131 (ST131) predominates globally among such MDR *E. coli* strains [11].

Our study was aimed at identifying the virulence and AMR genetic determinants predominant in MDR *E. coli*. We constructed core genome phylogeny using high quality SNP information to infer the genome wide association of virulence and resistance attributes among the sequenced samples.

## Materials and Methods

### Isolates and identification

A total of 99257 specimens were screened from BSI during the year 2015 to 2016 from the patients attending Christian Medical College, Vellore, India. Isolation and identification of the organism were carried out using a standard protocol [12]. Of the 1100 *E. coli* positives, 10% were observed resistant to carbapenems, of which 60 MDR isolates were selected in random for further characterization.

### Antimicrobial susceptibility testing (AST)

#### Disc diffusion

AST testing was carried out using the Kirby-Bauer disk diffusion method. The antimicrobial agents tested were Amikacin (30 μg), netilmicin (30 μg), gentamycin (10 μg), chloramphenicol (30 μg), ciprofloxacin (5 μg), cefotaxime (30 μg), cefoxitin (30 μg), ceftazidime (30 μg), cefpodoxime (10 μg), piperacilllin-tazobactam (100/10 μg), cefoperazone-sulbactam (75/30), imipenem (10 μg) and meropenem (10 μg), tigecycline (15 μg) and tetracycline (30 μg) according to guidelines suggested by CLSI M100-S27, 2017. Quality control strains (*K. pneumoniae* ATCC 700603, *P. aeruginosa* ATCC 27853 and *E. coli* ATCC 25922) were used in all batches as recommended by the Clinical and Laboratory Standards Institute.

#### Minimum Inhibitory Concentration (MIC) for colistin

Colistin MIC for the studied isolates was determined by broth microdilution and interpreted using CLSI 2017 breakpoint recommendations. *mcr-1* positive *E. coli* with the expected range 4 – 8 μg/ml, *E. coli* ATCC 25922 (0.25 – 2 μg/ml) and *P. aeruginosa* ATCC 27853 (0.5 – 4 μg/ml) were used as quality control (QC) strains for colistin MIC determination.

### Next generation sequencing and genome assembly

Genomic DNA was extracted using a QIAamp DNA Mini Kit (QIAGEN, Hilden, Germany). Whole genome sequencing (WGS) was performed using an Ion Torrent™ Personal Genome Machine™ (PGM) sequencer (Life Technologies, Carlsbad, CA) with 400-bp read chemistry according to the manufacturer’s instructions. Data were assembled with reference *E. coli* strain (NC000913) using Assembler SPAdes v.5.0.0.0 embedded in Torrent Suite Server v.5.0.3.

### Genome annotation

The assembled sequence was annotated using PATRIC, the bacterial bioinformatics database and analysis resource (http://www.patricbrc.org), and NCBI Prokaryotic Genomes Automatic Annotation Pipeline (PGAAP, http://www.ncbi.nlm.nih.gov/genomes/static/Pipeline.html). Downstream analysis was performed using the CGE server (http://www.cbs.dtu.dk/services) and PATRIC. The resistance gene profile was analysed using ResFinder 2.1 from the CGE server (https://cge.cbs.dtu.dk//services/ResFinder/). The sequences were also screened for antimicrobial resistance genes in the Antibiotic Resistance Genes Database (ARDB) and Comprehensive Antibiotic Resistance Database (CARD) through PATRIC. Virulence genes from the genomes were identified using VirulenceFinder 2.0 (https://cge.cbs.dtu.dk/services/VirulenceFinder/). Serotype of the isolates were identified using SerotypeFinder 1.1 (https://cge.cbs.dtu.dk/services/SerotypeFinder/).

### Genome based MLST analysis

Sequence types (STs) were analysed by MLST 1.8 (MultiLocus Sequence Typing) (https://cge.cbs.dtu.dk//services/MLST/). To visualize the possible evolutionary relationships between isolates, STs of the study isolates and the globally reported strains were computed using PHYLOViZ software v2.0 based on goeBURST algorithm. The study used Warwick database for all sequence based MLST analysis of *E. coli*.

### Genome comparison analyses

Gview, interactive genome viewer was used to compare the annotated *E. coli* genome arrangements with the reference *E. coli* K12 genome (NC_000913) [13]. Core genome analysis was performed using Roary: the Pan Genome Pipeline v3.11.2 from Sanger Institute [14]. Further, the tree file was visualised and analysed in iTOL v4 (https://itol.embl.de/).

This Whole Genome Shotgun project has been deposited at GenBank under the accession numbers PVPX00000000, PVPW00000000, PVPV00000000, PVPU00000000, PVPT00000000, PVPS00000000, PVPR00000000, PVPQ00000000, PVPP00000000, PVPO00000000, PVPN00000000, PVPM00000000, PVPL00000000, PVPK00000000, PVPJ00000000, PVPI00000000, PVPH00000000, PVOS00000000, PVPG00000000, PVPF00000000, PVPE00000000, PVPD00000000, PVPC00000000, PVPB00000000, PVPA00000000, PVOZ00000000, PVOY00000000, PVOX00000000, PVOW00000000, PVOV00000000, PVOT00000000, RCAC00000000, RCAE00000000, RCAI00000000, RCAH00000000, RCAJ00000000, RCAD00000000, RCAG00000000, RCAF00000000, RCAL00000000, RCAN00000000, RCAM00000000, RCAK00000000, SAZJ00000000, SAZP00000000, SAZU00000000, SAZV00000000, SAZF00000000, SAZG00000000, SAZH00000000, SAZI00000000, SAZK00000000, SAZL00000000, SAZM00000000, SAZN00000000, SAZO00000000, SAZQ00000000, SAZR00000000, SAZS00000000, SAZT00000000. The version described in this manuscript is version 1.

## Results

### Antimicrobial susceptibility

All 60 *E. coli* isolates were resistant to carbapenems, quinolones, cephalosporins and beta-lactamase inhibitors (Table S1). Whereas all the isolates were susceptible to colistin except B7532 and B9021, which exhibited an MIC of 32 μg/ml.

### Whole genome sequence analysis

#### Phylogeny of MDR E. coli

MLSTFinder revealed the different sequence types of the isolates. The study isolates belonged to 6 clonal complexes with 14 different sequence types. Few of the sequence types were observed to share same founder types revealing the evolution of these strains. CC10 and CC 405 were the two major CCs observed with ST-167, ST-410 and ST-405 as the common STs. Interestingly, nine isolates belonging to CC/ST-131 were identified, of which, all were of H-30 clade, except the isolate BA9313 (H-24). goeBURST analysis revealed the relation between sequence types observed within same clonal complex (Figure 1).

**Figure 1:**
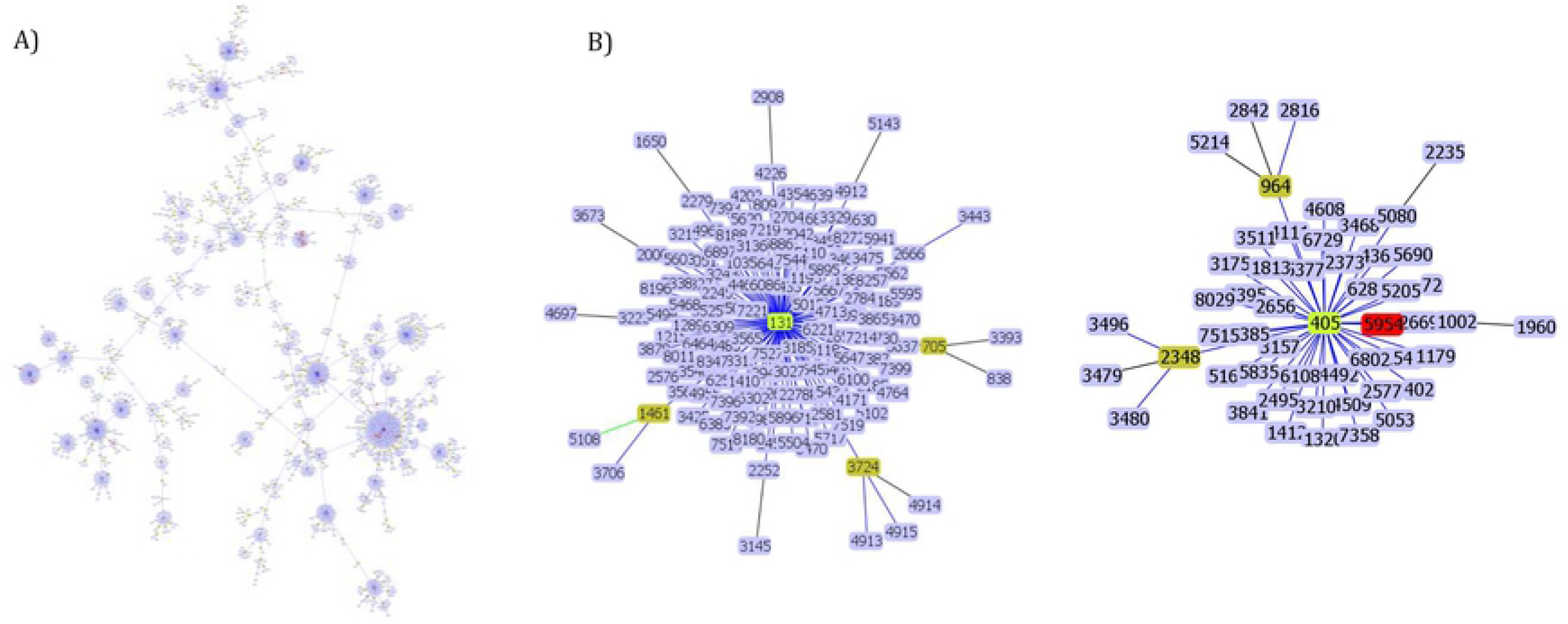
A) goeBURST image of *E. coli* showing clonal complex 10 with STs 44, 167, 410, 448, 617, 1702 and 2851, B) goeBURST image of *E. coli* showing clonal complex/ ST-131 (a), and CC405 (b) with STs 405 and 5954.

#### E. coli genome comparison

Whole genome composition of 60 MDR *E. coli* was compared with the *E. coli* K-12 reference genome which shoes the region of differences between these genomes (Figure S1). Totally 2518792 SNPs were identified in all analyzed genomes. On minimum 5957 and maximum 74713 SNPs were identified in each of the 60 MDR *E. coli* genomes when compared to the reference genome.

**Figure S1:** Circular genome plot comparing 60 MDR *E. coli* thereby showing differences in genome composition in comparison to the reference genome NC000913 *E. coli*.

### Core vs pan genome

Comparison between the core and pan genomes of 60 MDR *E. coli* isolates revealed 2258 core genes across all 60 isolates among the 17944 total gene clusters. This includes 600 soft core genes in 57 to 59 isolates, 3984 shell genes in 9 to 57 isolates and 11102 genes in less than 9 isolates (Figure 4). Further the pan genome matrix of the isolates depict the similarity between core genes and pan genes. Low similarity was observed across the genomes for the pan genes with high gene numbers and vice versa for the core genes (Figure 2).

**Figure 2:**
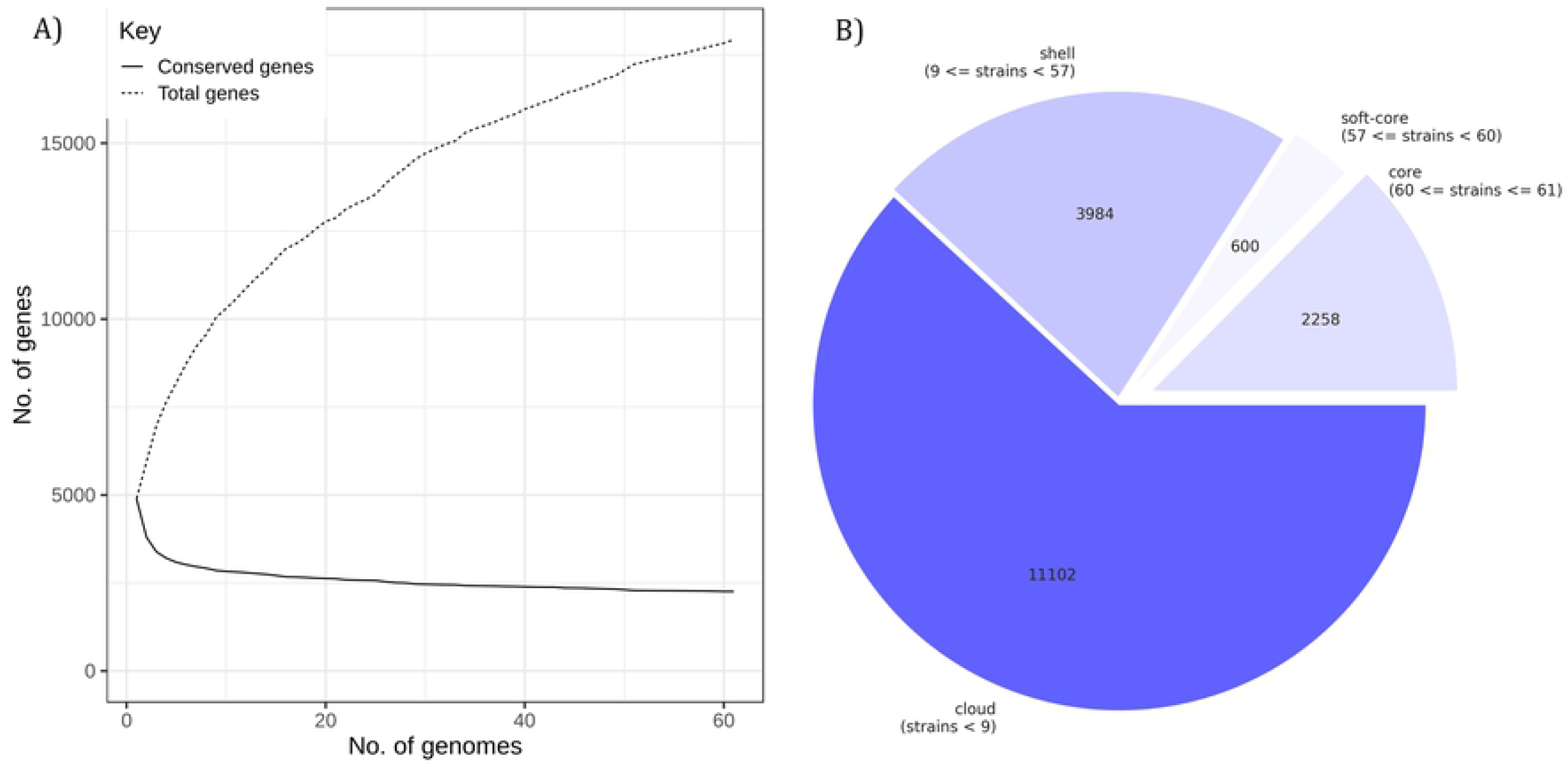
Pan genome vs core genome comparison depicting high number of pan genomes and low number of conserved genes A), Break-up of core genes, soft core genes and pan genes B) among the 60 MDR *E. coli* isolates from BSI

**Figure 3:**
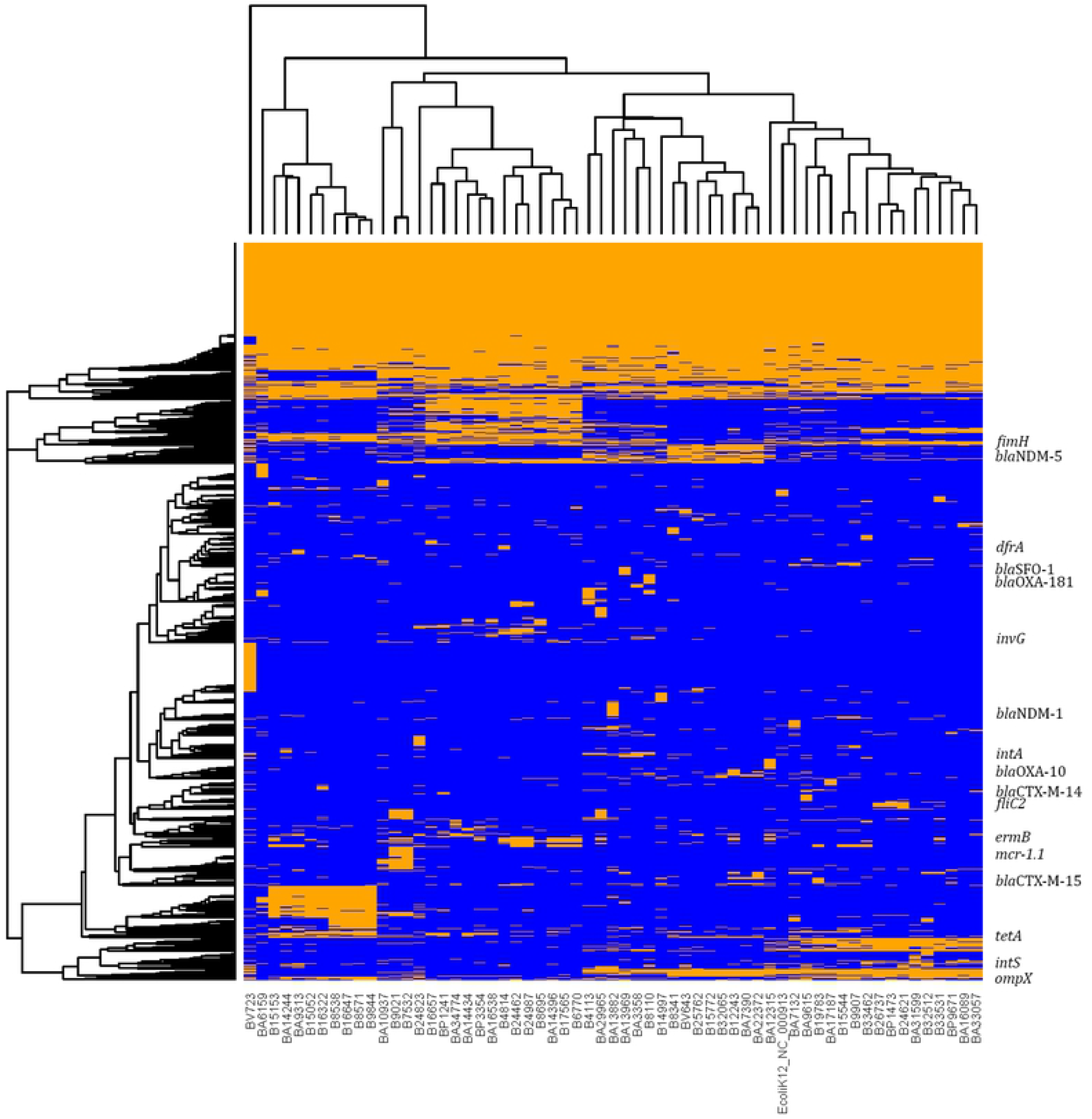
Pan genome matrix identified with the core gene sets of 60 MDR *E. coli* isolates depicting the important antimicrobial resistance (*bla*NDM-5, *bla*CTX-M-15, *bla*OXA-10, *mcr-1.1*) and virulence genes (invG, fimH, fliC2, intA) respectively.

**Figure 4:**
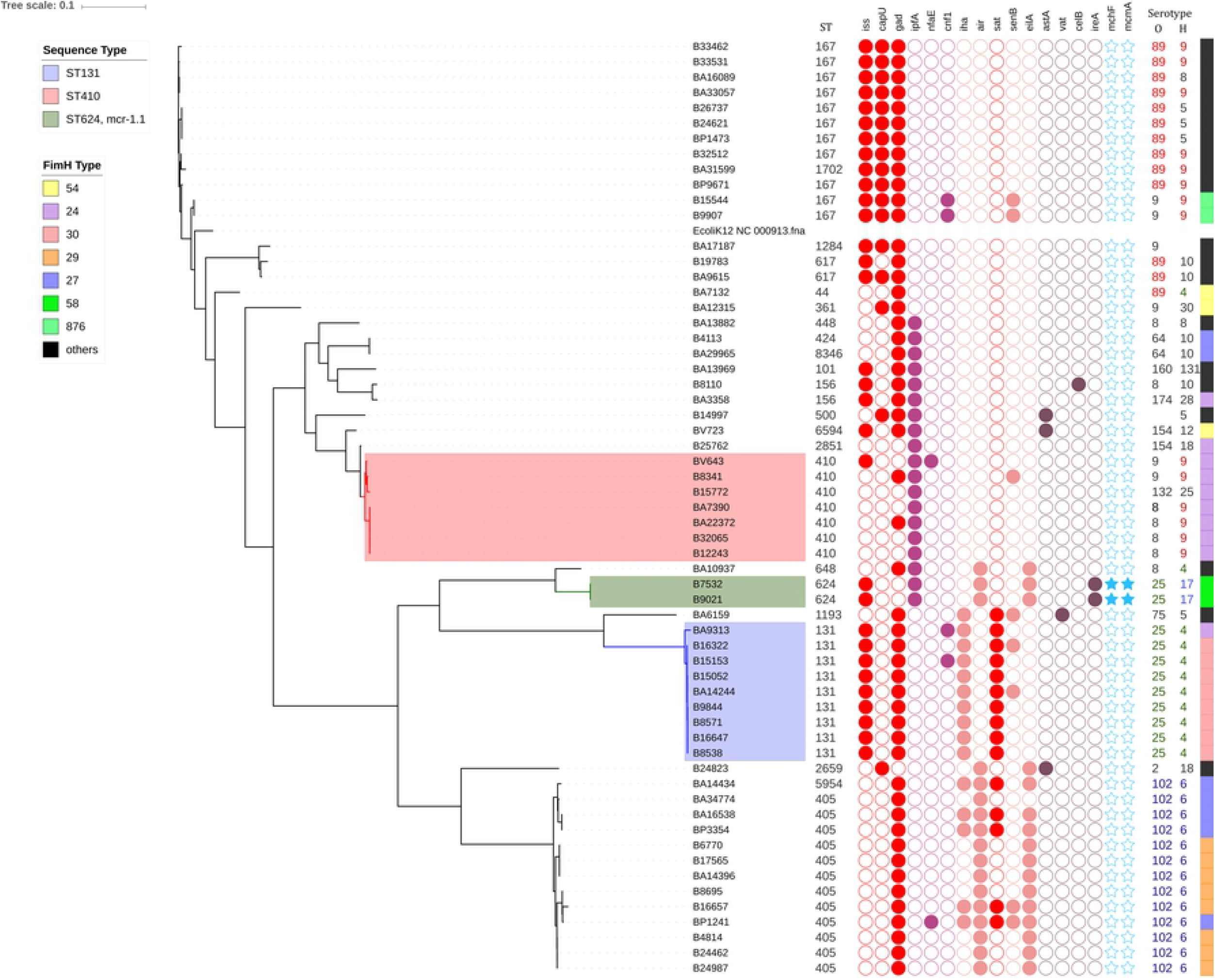
SNP phylogeny based comparison of genetic virulence traits observed in MDR *E. coli* strains exhibiting, sequence types, virulence gene profile, O and H antigens and Fim-H types.

**Figure 5:**
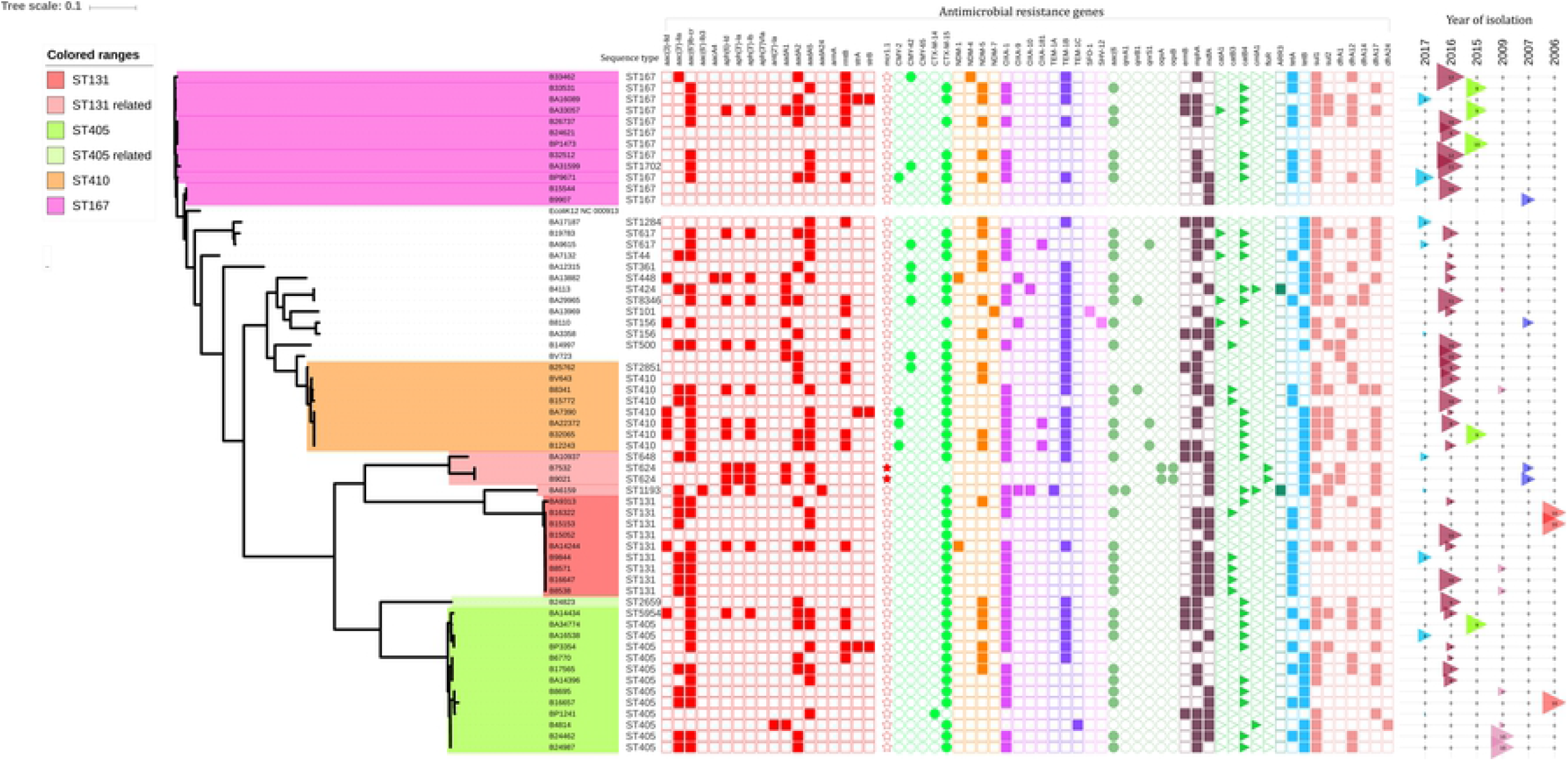
Antimicrobial resistance genes observed in MDR *E. coli* compared to the SNP based phylogeny. Depicting prevalence of *bla*NDM-5 among carbapenemases and *bla*CTX-M-15 among cephalosporinases.

#### Serotype prediction

Based on the whole genome data, serotype of the MDR *E. coli* isolates were detected. O102:H6 was the commonest serotype (18.75%) followed by O89:H9 (15.6%), O8:H9 (12.5%), O89:H5 (9.4%) and other serotypes. Few isolates had partial alignment of O or H antigen genes with the reference strains which further needs to be confirmed by additional tests.

#### Genetic virulence factors of MDR E. coli

Three common virulence gene profiles were observed among the isolates as follows, i) *iss, capU, gad*, ii) *ipfA*, and iii) *eilA, gad, air*. The FimH virulence typing revealed the types 5, 24, 27, 28, 30, 35, 54, 191, 54-like in comparison to the database.

#### Comparison of virulence and clonal traits

The virulence gene profiles of the 60 isolates were then compared to the FimH virulence types, MLST sequence type and SNP phylogeny. The *E. coli* isolates showed distinct groups including ST-131; H-30 clade based on the virulence genes identified (Figure 6). The sequence types were linked to the groups of same virulence gene profile and FimH virulence types.

#### Antimicrobial resistance genetic determinants

ResFinder revealed the presence of multiple AMR genes in each of the MDR *E. coli* (Figure 7). Aminoglycoside and beta lactam resistance genes were seen high compared to other groups. Among the aminoglycoside resistance genes, *aadA5* and *aac(6’)lb-cr* were higher followed by *aadA2* and *rmtB*. While among beta lactamases, *bla*CTX-M-15 was higher followed by *bla*NDM-5, *bla*OXA-1 and *bla*TEM-1B. Similarly, most of the isolates harboured *mphA, catB4, sul1, tetB, dfrA17* and *dfrA12*. Interestingly, two isolates, B7532 and B9021 carried *mcr-1.1* responsible for plasmid-mediated colistin resistance. The two isolates also showed phenotypic resistance to colistin with high MIC (>32 μg/ml).

## Discussion

The increasing usage of third-generation β-lactams and β-lactam inhibitors was accompanied with increases in prevalence of the MDR phenotype among *E. coli*. A study from India with invasive *E. coli* isolates between 2014 and 2016 reported susceptibility to cefoxitin 53%, ceftazidime 33%, cefotaxime 26%, ceftriaxone 25%, cefepime 29%, piperacillin tazobactam 66%, imipenem and meropenem 89%, aztreonam 36%, ciprofloxacin 19%, levofloxavin 23%, and amikacin 91% [15]. Among these, about 64% of *E. coli* were found to be ESBL producers.

The increasing frequency of antimicrobial resistance in clinical *E. coli* isolates was known to be associated with genetic determinants such as *bla*_CTX-M_, *bla*_NDM_, and *mcr* genes. In this study, multiple AMR genes for beta lactams, carbapenems, fluoroquinolones, tetracycline, aminoglycosides and colistin were identified. The presence of genotypic AMR genes correlated well with phenotypic expression for beta lactams, carbapenems, fluoroquinolones and tetracycline. Plasmids IncFII majorly carried AMR genes *bla*_CTX-M-15_, *bla*_NDM-5_, *aadA2, rmtB, sul1, drfA12, erm*(B) and *tetA*, while IncFI plasmids carried mostly *aadA5, sul2, dfrA17, mph*(A) and *tetB* genes [16].

The MDR *E. coli* isolates upon SNP based phylogeny grouped to four major clades, ST167, ST410, ST405 and ST131. Among the isolates, for carbapenem resistance, *bla*_NDM-5_ was common in ST131, ST405 and ST410 clades. While among ST167, *bla*_NDM-4, −5_ and_-7_ were seen. Whereas, previous reports identified *bla*_NDM-1_ as common among *E. coli* though the sample size was lesser [17], whereas in China, *bla*_NDM-1_ and *bla*_NDM-5_ were seen in equal numbers [18].

*bla*_OXA-48_ type carbapenemases are the most commonly reported in *E. coli* [19] followed by *bla*_NDM_ [20], *bla*_IMP_ [21] and *bla*_KPC_ [22] type carbapenemases have also been determined. Studies report occurrence of *bla*_OXA-48_ from as low as 3% to 22% [23, 9]. A report from India has shown, *bla*_NDM_ was common among carbapenemases in *E. coli* (70%), followed by *bla*_OXA-48_ (24%) and *bla*_VIM_ (17%). *bla*_NDM_ alone is 48%, *bla*_OXA-48_ alone is 19%, *bla*_NDM_+*bla*_OXA-48_ is 5%, *bla*_NDM_+*bla*_VIM_ is 17% [24]. In our study, the combinations observed are *bla*_NDM-1_ - 1, *bla*_NDM-4_ - 1 *bla*_NDM-5_ – 6 *bla*_NDM-7_ – 1, *bla*_OXA-1_ - 21, *bla*_OXA-181_ - 2, *bla*_NDM-5_+*bla*_OXA_ - 16, *bla*_NDM-5_+*bla*_OXA-181_ – 1, *bla*_NDM-1_+*bla*_OXA_ – 2. This revealed that *bla*_OXA-1_ was predominant followed by *bla*_NDM_ in carbapenem resistant *E. coli* in our setting, though *bla*_OXA-181_ was rare.

In addition, aminoglycoside resistance had been an important concern among Gram-negatives. Acquired 16S-RMTases were known to confer extremely high level of aminoglycoside resistance, due to which key aminoglycosides including gentamicin, tobramycin, and amikacin are ineffective against carbapenem resistant strains [25].

Accordingly, plazomicin, a new aminoglycoside agent identified to combat against carbapenem-resistant Enterobacteriaceae, was found inactive if the isolates co-produced 16S-RMTases [26]. In this study, about 90% of the RMTase positive *E. coli* co-harbored carbapenemases, which further adds to the burden of carbapenem resistance. Similarly, Taylor et al. [27] and Poirel et al. [28] had reported 83.1% and 45.4% co-occurrence of carbapenemases in 16S RMTase producing Enterobacteriaceae, respectively.

For cephalosporin resistance, the isolates of ST131 and ST405 clades carried *bla*CTX-M-15, whereas ST167 and ST410 carried *bla*CTX-M-15 and *bla*_CMY_ genes. Previously 54.34% *bla*_CTX-M_ was reported in ESBL positive isolates from India [29]. Recently, plasmid-mediated colistin resistance is increasingly reported from *E. coli* [30–32]. This study observed two isolates (B7532, B9021) with *mcr-1.1* expressing high MIC of >32 μg/ml to colistin. Though the isolates were from 2007, no records of *mcr* in *E. coli* were observed after 2007.

The antimicrobial susceptibility of *E. coli* has been shown to vary geographically [33]. Among the different clonal groups observed, *E. coli* ST131 was most commonly associated with community acquired infection, which recently were highly associated with healthcare settings. ST131 was reported as the predominant lineage carrying *bla*_CTX-M-15_ and other ESBLs. Most of the MDR *E. coli* carrying *bla*_CTX-M-15_ from different countries in Europe and North America were homogenously grouped into the *E. coli* O25:H4-ST131 [6,35–36]. In this study, 87% of the isolates carried *bla*_CTX-M-15_, among various STs, where only nine isolates belong to ST131. Among the observed STs in this study, *bla*_CTX-M-15_ was previously reported for its strong association among ST167, ST617, ST405, ST410, ST131 and ST361 [36].

Though ST131 clones were predominantly reported worldwide, the STs observed in this study were known for its distinct lineages. ST167 was previously reported for its ability to carry *bla*_NDM_ genes in China [37–39]. ST405 had been known as another global clonal group similar to ST131 [40]. ST405 lineage was reported to carry *bla*_NDM_ genes in hospital settings [41] and also reported to carry blaKPC-2 [42]. In addition, ST405 was reported as a lineage carrying fluoroquinolone resistance in Japan [40]. Recently, ST410 was reported as a possible international high risk clone with B2/H24R, B3/H24Rx, and B4/H24RxC AMR clades. B3/H24Rx was reported to be evolved by acquisition of the *bla*_CTX-M-15_ and an IncFII plasmid. B4/H24RxC emerged by acquiring IncX3 plasmid with *bla*_OXA-181_ known for carbapenem resistance, which further acquired *bla*_NDM-5_, on a conserved IncFII plasmid [43]. In this study, all ST410 isolates (*n* = 7) harboured *bla*_CTX-M-15_ gene, while only B25762, BV643, B12243 and B32605 had IncFII plasmids and *bla*_NDM-5_ (B3/H24RxC). B12243, in addition harboured IncX3 with *bla*_OXA-181_ (B4/H24RxC), while BA22372 and BA9615 had only *bla*_OXA-181_ in IncX3 plasmid (B4/H24RxC).

Virulence genes observed among the *E. coli* isolates varied according to the different clades observed. The comparison of the virulence gene type with SNP based phylogeny revealed the acquaintance and deletion of virulence genes. Genes *iss, capU* and *gad* were observed in ST167 clade. ST131 possessed *iha, sat, cnfl* and *senB* in addition to *iss* and *gad*. ST131 had lost the *capU* genes. Further, ST405 clade also lost *iss* and gained *eilA* and *air* genes with FimH type 29. Few isolates of ST405 retained *iha* and *sat* genes belonging to FimH 27 type within ST405. This was followed by ST410 (FimH 24) that predominantly had *ipfA* gene and lost all other genes, except few. Overall, gad gene served as backbone for ST167, 131 and ST405 clades, while *ipfA* for ST410. Similar studies comparing the evolution of virulence pattern with phylogeny is still lacking.

FimH had been reported as a major candidate for the development of a vaccine against pathogenic *E. coli* [44] which was responsible for mannose-sensitive bacterial adhesion [45]. Though high nucleotide conservation of >98% was observed in *fimH* alleles, minor sequence differences were reported to correlate with differential binding and adhesion phenotypes [45].

Fim-H types correlated with the STs observed in the study isolates. FimH 27 and 29 type grouped with ST405 clade. The ST405 clade with 27 and 29 fimH types would have possibly emerged from the ST131 clade with Fim-H type 24 and 30. However, another clade with FimH 24 belonged to ST410 with CC10. Similar study comparing the MLST and fim-H types were previously reported by Dreux et al. [46] in Adherent-Invasive *E. coli*.

Moreover, concatenated SNPs based phylogeny revealed higher discrimination between the clinical MDR *E. coli* isolates and disclosed the evolutionary pattern with accumulated SNPs. Among the study isolates, even within the clones, evidence of evolution is seen with the difference in root to tip. This suggests continuous evolution leading to diversified MDR *E. coli* strains in India.

## Conclusions

To the best of our knowledge, this is the first report on SNP phylogeny in comparison with AMR and virulence traits in *E. coli* in India. The study revealed the prevalence of *bla*_NDM-5_ among the clades ST131, ST405 and ST410 clades. *bla*_CTX-M-15_ was responsible for cephalosporin resistance in ST131 and ST405 clades whereas, ST167 and ST410 carried both *bla*_CTX-M-15_ and *bla*_CMY_ genes. For aminoglycoside resistance, *rmtB* was positive for 31.7% of the isolates, of which 30% were co-harboring carbapenemases. The FimH types and virulence gene profile positively correlated with the SNP based phylogeny. However the predominant ST131 epidemic clone was smaller the study population while ST167 and ST405 clones with multiple AMR genes were predominant. Isolates with *iss, capU* and *gad* virulence genes were the major type. Moreover, SNP based phylogeny revealed the evolution of the MDR clones among the study population with the accumulation of SNPs, which suggests continuous molecular surveillance to understand the spread of MDR clones in India.

